# Online stimulation of prefrontal cortex during practice increases motor variability and modulates later cognitive transfer: a randomized, double-blinded & sham-controlled tDCS study

**DOI:** 10.1101/2023.12.22.572904

**Authors:** Nisha Maria Prabhu, Nico Lehmann, Elisabeth Kaminski, Notger Müller, Marco Taubert

## Abstract

**Background:** The benefits of learning a motor skill extend to improved task-specific cognitive abilities. The mechanistic underpinnings of this motor-cognition relationship potentially rely on overlapping neural resources involved in both processes, an assumption lacking causal evidence.

**Objectives:** We hypothesize that interfering with prefrontal networks would affect concurrent motor skill performance, long-term learning and associated cognitive functions dependent on similar networks (transfer).

**Methods:** We conducted a randomized, double-blinded, sham-controlled brain stimulation study using transcranial direct current stimulation (tDCS) in young adults spanning over three weeks to assess the role of the prefrontal regions in learning a complex balance task and long-term cognitive performance.

**Results:** Balance training combined with active tDCS led to higher performance variability in the trained task as compared to the sham group, without affecting the learning rate. Furthermore, active tDCS also positively impacted performance in untrained motor and cognitive tasks.

**Conclusion:** The findings of this study help ascertaining the networks directly involved in learning a complex motor task and its implications on cognitive function. Hence, opening up the possibility of harnessing the observed frontal networks involved in resource mobilization in instances of aging, brain lesion/injury or dysfunction.

## 1. Introduction

Physical activity has proven instrumental in enhancing overall health and well-being across the lifespan. Physical inactivity on the other hand, particularly in the context of aging, has dire consequences extending to cognitive dysfunction [1,2]. Gaining a better understanding of the effects of physical activity on cognitive performance is crucial to support healthy aging, or ameliorate cognitive impairments by incorporating spared mobility into therapy. Perhaps a key component in explaining the link between physical activity and cognition lies within the brain and the synergistic neural networks subserving both motor processing and cognitive functions. Although colocalized brain activity has been identified for motor and cognitive processes, we lack important causal evidence linking movement and cognition at the neural level [3,4].

Among wide-ranging forms of physical exercise, the influence of complex motor skill learning on cognition has garnered considerable attention. Skill learning has the potential to positively impact cognition through the involvement of key cognitive functions supporting learning, viz., by challenging functions like information processing, decision-making, movement selection, planning, exploring-tracking-switching between courses of actions and predicting outcomes based on experience [5]. While the prefrontal cortex (PFC) has been associated with the majority of the above-mentioned cognitive functions [6–9], the PFC is also capable of undergoing motor learning-induced brain plasticity. For example, learning a complex and challenging whole-body task (DBT-dynamic balance task) was shown to induce structural and functional changes in the PFC, with PFC structure predicting improved balance task learning [10–12]. Moreover, various training studies suggest transfer effects of motor balance training to relevant cognitive domains [13–15]. The neural overlap hypothesis predicts that behavioural transfer from motor practice to cognitive performance is sub-served by overlapping neural circuits [16–18]; and its underlying mechanisms are hypothesized to occur during the acquisition period of a new skill [19]. Despite these observational neuroimaging and behavioural findings, the causal role of the PFC in motor balance learning and its potential to mediate learning-induced cognitive transfer remains unclear.

Unravelling complex brain-behaviour relationships has been effectively achieved through non-invasive brain stimulation techniques (NIBS). Transcranial direct current stimulation (tDCS) is one stimulation technique widely employed in the context of motor learning. tDCS involves modulating cortical excitability of a target brain region [20]. Anodal tDCS over the primary motor cortex (M1) was shown to enhance motor learning [21,22]. Improved overnight motor skill consolidation has been observed through an effect on networks involved in early consolidation after anodal tDCS over M1 [23]. When learning is driven by factors such as performance feedback, sensory feedback error signals and cognitive strategies, opportunity for variation within the learning process is created in an attempt to explore the solution space [24–26]. In such cases, tDCS over PFC has resulted in faster motor learning through regulation of performance variability [27]. tDCS thereby aids in deriving causal inferences in the face of correlative electrophysiological or neuroimaging evidence [28,29]. Although the neurophysiological effects of a single tDCS session are shown to last a few minutes to a couple of hours after the end of stimulation [30,31], it is nevertheless capable of inducing long-term structural plasticity in the form of rearranged synaptic networks and spinogenesis as established through animal models [32,33]. Similarly, in older adults, training combined with tDCS spread over multiple days was shown to modulate functional connectivity and microstructural brain alterations associated with cognitive performance gains [34]. In line with these findings, prefrontal tDCS applied during motor practice may therefore influence the balance learning-induced prefrontal neural changes that support ongoing balance performance; also affecting the transfer to cognitive tasks assessed after a time delay that outlasts the acute neurophysiological effects of tDCS.

In order to test this prediction, cathodal tDCS (c-tDCS) over right PFC (rPFC) [10,35] was used during DBT practice sessions. First, we hypothesized cathodal compared to sham-tDCS will affect performance indices and learning of the DBT task. We have shown prefrontal regions to undergo structural changes throughout long-term DBT practice [11] and rapid grey matter changes in M1 after a single DBT practice session [36]. Therefore, we aim to assess the role of PFC during the process of skill learning using concurrent tDCS over several training sessions. Following the neural overlap hypothesis, we further predict prefrontal tDCS during motor practice to modulate remote (24h after motor practice) performance in cognitive tasks that rely on overlapping prefrontal networks without affecting cognitive performance immediately after stimulation.

## 2. Material and methods

### 2.1. Ethics statement

This study was approved by the local ethics committee of the Otto-von-Guericke University, Magdeburg [130/20]. Conforming to the declaration of Helsinki, all subjects provided their written informed consent prior to participation in the experiment and received financial compensation for participation.

### 2.2. Study design

We conducted a randomized, double-blinded, sham-controlled study to examine the modulatory effect of c-tDCS over the PFC during balance performance and learning over 3 weeks in forty-four subjects between the ages of 18-35yrs (n=44, 21.8±3.25yrs, 27 females). Sample size was estimated based on findings from [35] using a similar motor learning paradigm along with concurrent tDCS (supplementary materials 1.1. for further details). Highly skilled subjects such as slackliners or participants with prior experience with the DBT were excluded. Additionally, in order to evaluate their general physical activity levels, participants were required to fill-in an activity questionnaire [37].

All participants were informed about potential risks of non-invasive brain stimulation used in this study. After granting their written informed consent, participants were randomly assigned to either cathodal (c-tDCS) or sham (s-tDCS) groups by one of the authors (MT: no contact with any of the participants). Neither the researchers involved in data acquisition/training nor the participants were aware of the group assignment. Irrespective of the training groups, similar tDCS electrode montage using EEG 10-20 position was applied. The entire training duration lasted a total of 3 weeks consisting of two training sessions per week (TD1-TD6) with motor and cognitive transfer tests conducted 24 hrs pre- and post-the training period (Figure 1). The first training session of the week (TD1, TD3, TD5) included DBT practice with concurrent c-tDCS or s-tDCS over right PFC (rPFC). These training sessions were followed (24hrs later) by a re-evaluation of the DBT performance without c-tDCS (TD2, TD4, TD6). To control for the acute effects of tDCS on general balance ability and general cognitive abilities of the participants, balance and cognitive assessments were performed as control tasks immediately before and after c-tDCS application (refer 2.2.3).

**Figure 1.**
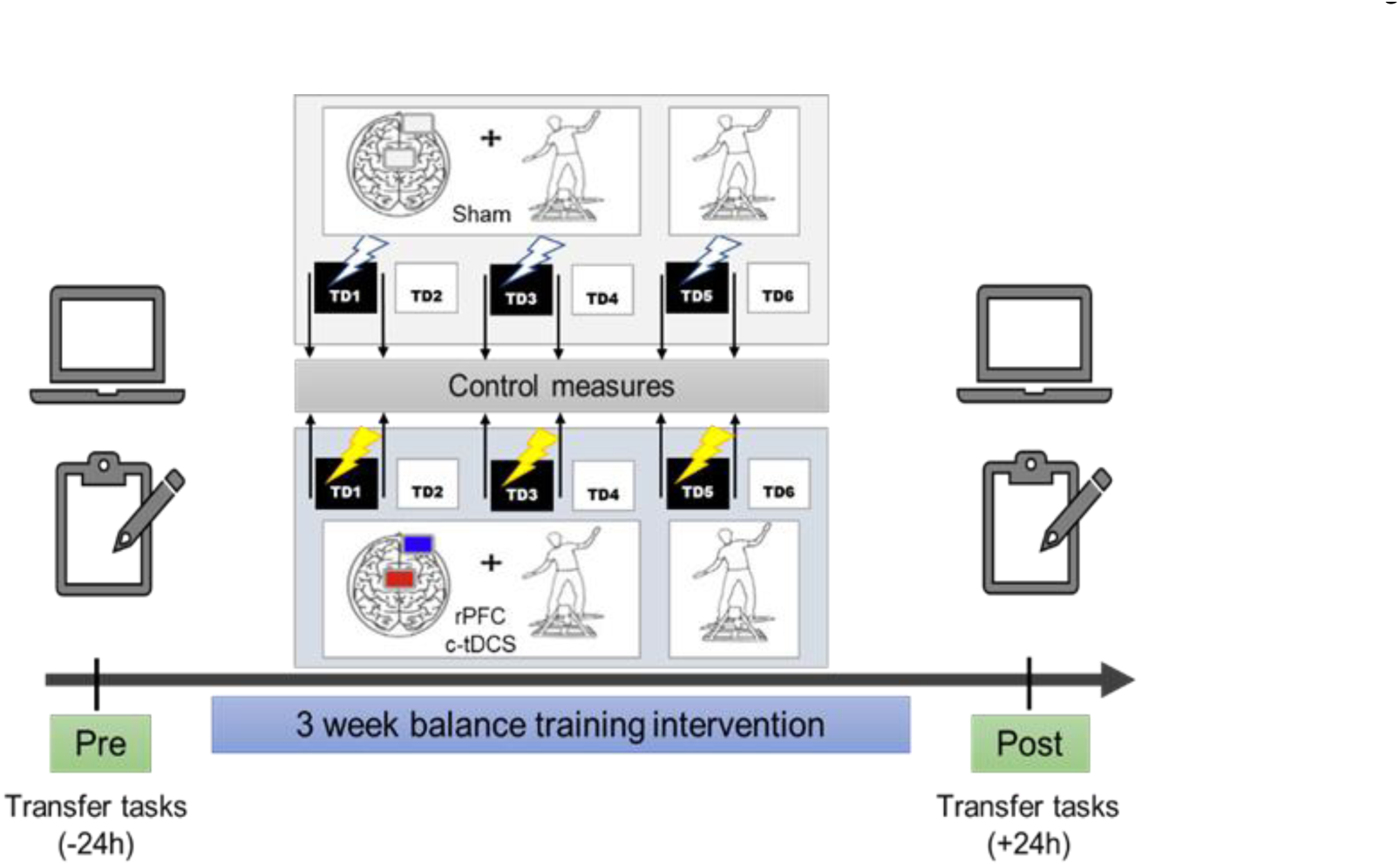
Experimental design: Participants trained on the DBT over 3 weeks with two practice sessions per week. The first session of the week included practice under tDCS stimulation followed 24 hours by practice without tDCS. Every session included 15 trials lasting 30 seconds each, interspersed with a rest period of 90 seconds. All participants also performed a battery of motor transfer, computer-based and paper-pencil cognitive tests before and after the 6 training sessions.

#### 2.2.1. Complex balance task (DBT)

The motor learning paradigm in our study included a whole-body dynamic balance task consisting of a balance platform that moves in a see-saw like manner known as a Stabilometer (stability platform, Model 16030, Lafayette Instruments, Lafayette, IN, USA), with a maximum deviation of 26 degrees on each side. A typical training session on the stabilometer included 15 trials lasting 30 secs each, with 90 seconds rest period between each trial. The goal was to maintain the platform in a horizontal position, i.e., parallel to the floor, for as long as possible during the 30sec trial; staying within a target deviation of 0°-3° to the right or left from the horizontal axis. This required the participant to position the body’s centre-of-pressure vertically above the boards’ axis of rotation. Each training session lasted approximately 30-40 mins each day. At the end of each trial, participants received feedback about their performance in the form of time in balance (TIB-outcome measure), i.e., seconds spent within the ±3° target window. Receiving no instructions regarding task performance strategies, apart from the necessary safety guidelines and TIB feedback, they were granted the freedom to explore their own strategies in order to improve performance over the 6 training sessions (Discovery learning approach)[38,39].

#### 2.2.2. Transcranial direct current stimulation (tDCS)

A weak direct current of 1mA generated from a rechargeable battery driven stimulator (NeuroConn Gmbh, Ilmenau, Germany) was used for a total duration of 20min during TD1, TD3, TD5. Electrodes were fastened using Velcro straps over the areas corresponding with rPFC (EEG 10-20 electrode placement), i.e., cathodal electrode on the right supraorbital region (Fp2). The reference electrode was placed midway between frontal and central zero (Fz-Cz-with slight off-set to left side) ensuring no overlap with the cathodal electrode occurred while simultaneously avoiding stimulation over the M1 area [21]. Electrodes were encased within sponge covers drenched in saline solution (NaCl) and rehydrated intermittently if necessary using syringes without moving the electrodes from their fastened position. Sizes of both electrodes were kept at 35cm^2^ (5×7cm) with a current density of 0.028 mA/cm^2^ and a total charge of 0.033 C/cm^2^ under each electrode, similar to [35]. The cathodal stimulation group (c-tDCS, n=22) experienced stimulation with a trapezoidal pulse form consisting of ramp-up at the beginning and ramp-down lasting 30 secs at the end of 20-min stimulation period. However, the s-tDCS group (n=22) received a similar ramp-like stimulation with a fade-in, maintenance of stimulation for 30 secs only, followed by a fade-out. The tDCS stimulation was started only after the second trial during each training session and lasted 20 minutes thereafter. The participants carried the stimulator in a backpack during DBT practice. As a precautionary measure, a questionnaire pertaining to sensory perception, changes in attention, perception of fatigue and discomfort after/ during stimulation was administered [40]. To assess the success of blinding, all participants were asked whether they believed they received stimulation or not after TD1, TD3 and TD5.

#### 2.2.3. Control measures

Acute effects of tDCS stimulation on general balance ability and executive functions were tested using the Balance Error Scoring System (BESS)[41] and the Stroop test [42,43] respectively. These tests were administered pre- and post-training sessions where participants received tDCS (refer supplementary materials 1.3 for test description). These tasks were chosen to match our tasks of interest with respect to its characteristics and difficulty, although distinct in terms of the involved cognitive or motor functions of interest. This allowed us to ascertain task specificity while examining the acute effects of tDCS, in turn avoiding confounds via co-affected supporting functions [28].

#### 2.2.4. Transfer tests

Based on transfer effects reported in previous coordinative exercise training studies with and without tDCS[4,14,18,44], a cognitive test battery conducted 24 hours before and after the training period investigated the transfer effects of concurrent tDCS and motor practice. The tests and the measured parameters included: Visual and Verbal Memory Test-delayed recall and rate of forgetting (VVM)[45], d2-Test of Attention-concentration score (d2-R)[46], Eriksen Flanker task-accuracy and reaction time interference [47] and Trail making test (TMT)-time to completion in TMT-A (1-2-3-…), TMT-B (1-A-2-B-…) and Δ TMT (factoring out the time component of TMT while accounting for completion times in both subtests TMT A & B)[48,49]. As a motor transfer test, a football header task available on the WiiFit console (Nintendo) was used to assess the goal-directed control of COM movement at the beginners and advanced level. (Detailed description of all the tests conducted are included in supplementary materials 1.2).

### 2.3. Data analysis

All statistical analyses for this study were conducted using the software R version 4.1.3 [50]. Between group comparisons at baseline for all demographic variables were conducted, depending on the scale level, using chi square or Brunner-Munzel [51] tests. To investigate the performance changes over the entire training duration and on the data from the questionnaire inspecting perceived sensory effects of stimulation, robust two-way mixed ANOVAs based on 20% trimmed means as implemented in the WRS2 package [52] were used. The blinding responses were analysed using BI package implemented in R. James blinding index (two-sided) for TD1, TD3 and TD5 are reported separately and interpreted as 0.0 = complete unblinding, 0.5 = random guessing and 1.0 = complete blinding [53].

The TIB recorded during 15 trials was averaged for each TD for between and with-in group comparisons. In addition to TIB, coefficient of variation (CoV= SD/mean) in TIB over each TD was compared. A posthoc analyses of CoV’s at every TD was further conducted using the nonparametric combination (NPC) framework, combining results from multiple studentized Wilcoxon permutation tests [51,54] using Fisher’s chi-square [55] combination into a single global *p*-value accounting for the dependence among the component tests (R package NPC v1.1.0)[56].

In order to investigate the effect of c-tDCS on learning-induced transfer, pre-post difference scores were calculated for (1) the control tasks, (2) cognitive and (3) motor transfer tasks. Transfer task comparisons were conducted using a non-parametric Brunner-Munzel test (brunnermunzel package)[51], whereas mixed ANOVAs (described above) were used for comparing control tasks accounting for multiple time points. Type I error rate α was set at the conventional significance level of .05. Depending on the statistical test used, effect sizes are reported as Cohen’s *d* (small = 0.20, medium = 0.50, large = 0.80) or Cliff’s delta (δ; Cliff, 1996) interpreted as small = 0.11; medium = 0.28; large = 0.43 [58].

## 3. Results

Baseline characteristics of the participants in this study with respect to age, gender, body height, body weight, hand dominance and day-to-day physical activities did not differ between groups (Table 1).

**Table 1.** Demographic data: Comparisons between groups in relation to age, gender, height, weight, dominance, physical activity. Values displayed denote the median and interquartile ranges within parentheses for both groups. All statistical comparisons performed with Brunner-Munzel test except gender and hand dominance (chi-square Χ^2^).

| Characteristics | Overall | c-tDCS | s-tDCS | c-tDCS vs s-tDCS |
| --- | --- | --- | --- | --- |
| Age | 21 (3) | 21 (2.75) | 21 (2.75) | $t(41.83) = 0.41, p = .68$ |
| Sex (F; M) | 27;17 | 13;9 | 14;8 | $X^2(1) = 0.09, p = .76$ |
| Height | 174 (15) | 173 (16.5) | 174.5 (12) | $t(39.87) = -0.33, p = .74$ |
| Weight | 69 (12.75) | 68 (16) | 70.5 (10.75) | $t(40.68) = -0.85, p = .39$ |
| Hand dominance (Left; Right) | 2;42 | 0;22 | 2;20 | $X^2(1) = 2.09, p = .15$ |
| Physical activity: Work index | 2.19 (1.63) | 2.13 (1.31) | 2.25 (1.38) | $t(40.13) = -0.29, p = .77$ |
| Physical activity: Sport index | 3.25 (0.88) | 3.13 (1.00) | 3.25 (1.00) | $t(41.90) = -0.78, p = .44$ |
| Physical activity: Leisure time index | 3.5 (0.75) | 3.5 (0.69) | 3.75 (0.63) | $t(40.09) = -0.76, p = .45$ |

### 3.1. Control measures

#### 3.1.1. Stimulation Questionnaire

In the questionnaire related to tDCS-induced immediate effects, participants rated on a scale from 0 to 4 their subjective perception of pain, attention, sensation, etc. No significant main or interaction effects were found on factors like tingling (*F*(2, 22.42) = 0.17, *p* = .84), burning sensation (*F*(2, 19.66) = 0.15, *p* = .85), headache (*F*(2, 22.63) = 1.18, *p* = .32), concentration problems (*F*(2, 21.15) = 0.58, *p* = .57), attention (*F*(2, 17.33) = 0.23, *p* = .79), etc. (refer supplementary material 2.1. for further details)

#### 3.1.2. Blinding of stimulation

The blinding index (BI) on TD1 was estimated at 0.56 with 95% CI [0.42, 0.69], on TD3 BI = 0.44 with 95 % CI [0.32, 0.57] and on TD5 BI = 0.59 with 95% CI [0.49, 0.69], indicating random guessing (Figure 2). These results combined with the results of the stimulation questionnaire, indicate successful blinding between the groups.

**Figure 2.**
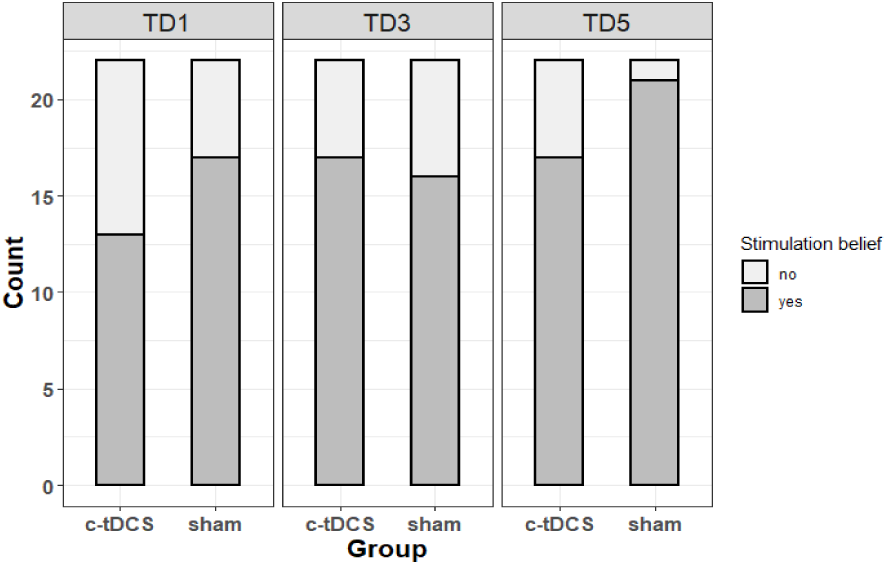
Responses from both groups about stimulation belief (blinding) on training sessions 1, 3, 5

#### 3.1.3. Stroop task

No significant effect for group, *F*(1, 25.31) = 0.33, *p* = .57, time *F*(2, 22.44) = 0.06, *p* = .95, or interaction effect, *F*(2, 22.44) = 1.90, *p* = .17, was detected for Stroop accuracy interference reduction. A significant main effect of group was found only for reaction time during the Stroop task, *F*(1, 25.32) = 7.53, *p* = .01, without an effect of time, *F*(2, 22.77) = 0.35, *p* = .71, or interaction, *F*(2, 22.77) = 0.14, *p* = .87 (Supplementary Figure. S7). This result stems from the poorer performance of the s-tDCS group immediately after training compared to pre-training performance (c-tDCS group performance remained unchanged). This pattern remained consistent over time.

#### 3.1.4. BESS task

tDCS stimulation did not affect general balance ability between groups as no effect for factor group (*F*(1, 25.96) = 0.71, *p* = .40), time (*F*(2, 22.04) = 1.18, *p* = .33) nor an interaction effect (*F*(2, 22.04) = 1.58, *p* = .23) was observed (Supplementary Figure. S6).

### 3.2. tDCS effects on DBT performance

#### 3.2.1. DBT learning

Baseline performance recorded as the first two trials on TD1 (before tDCS stimulation commenced) was found to be similar between both groups (mean TIB c-tDCS: 3.05 ± 1.7 secs vs s-tDCS: 2.99 ± 1.49 secs), Brunner-Munzel *t*(41.97) = -0.27, *p* = .78, δ = .05 (Supplementary Figure. S1). After six consecutive training sessions on the stabilometer, both groups significantly improved their DBT performance, *F*(5,18.66) = 34.57, *p* = .00, *d* > 1.81 (mean TIB c-tDCS: 13.53 ± 4.5 secs and s-tDCS: 13.36 ± 3.93), without an effect of group (*F*(1,25.55) = 0.13, *p* = .73, *d* = .11) or group*time interaction effects, *F*(5,18.66) =1.29, *p* = 0.31, *d* = .35 (Figure 3A).

**Figure 3.**
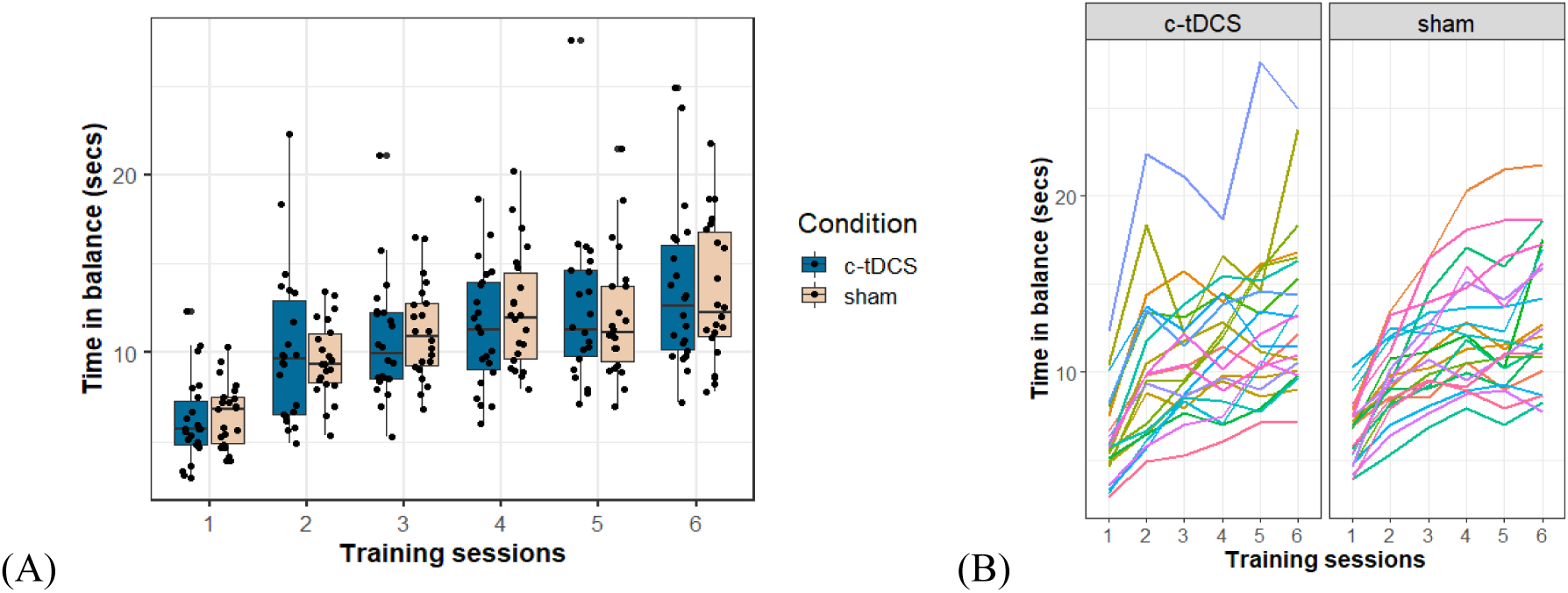
DBT performance and learning: (A) Improvements in Time in Balance (TIB) from training day-1 to training day-6. Every data point represents each participants TIB values; (B) Trajectory of performance change for every participant over six training sessions in the c-tDCS and s-tDCS groups, respectively. Each line represents the performance trajectory of a different participant.

#### 3.2.2. DBT performance variability

The c-tDCS group (0.24 ± 0.03) on average exhibited significantly larger performance variability compared to s-tDCS (0.21 ± 0.025), Brunner Munzel *t*(38.16) = -2.22, *p* = .03, δ = .36 (medium)(Figure 4A). Across the six training sessions (Figure 4B), the c-tDCS group displayed higher CoV than the s-tDCS group, *F*(1,22.79) = 4.91, *p* = .04, *d* = .68. CoV reduced significantly over time for both groups *F*(5,19.36) = 52.07, *p* = .00, *d* > 2.23, revealing a group*time interaction *F*(5,19.36) = 2.96, *p* = .04, *d* = .53

**Figure 4.**
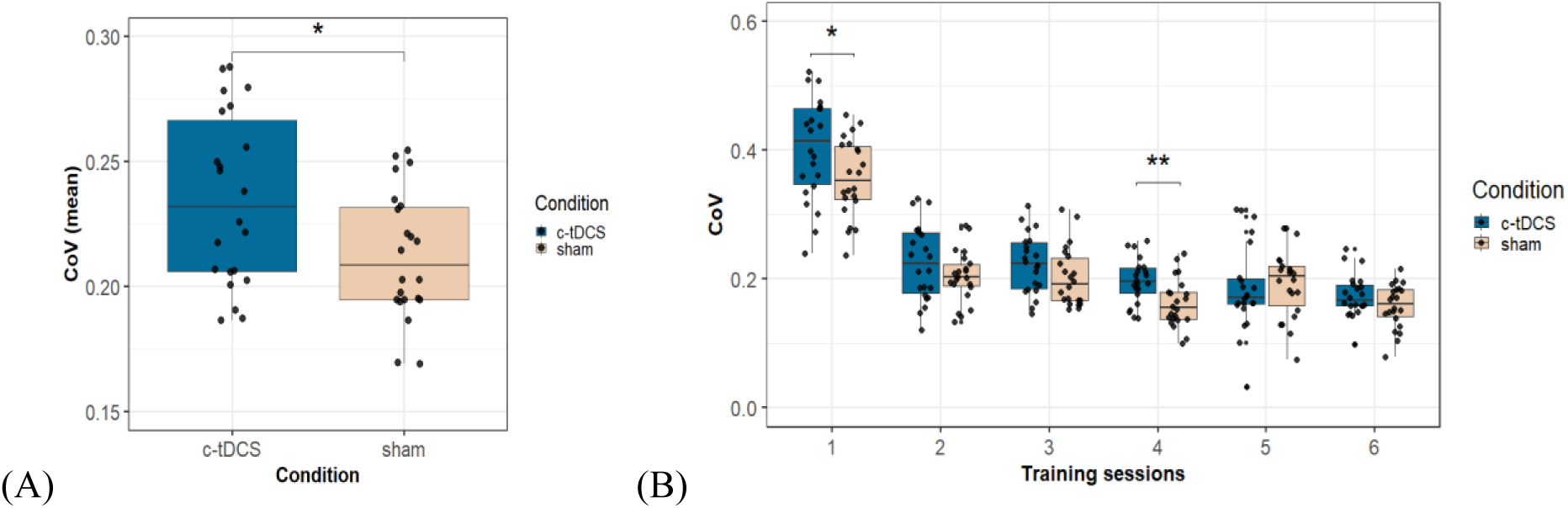
Motor variability expressed as coefficient of variation (CoV). (A) Mean CoV over the entire training duration shows significantly higher variability exhibited by the c-tDCS group, Brunner-Munzel t(38.16) = -2.22, p = .03, δ = .36 (medium); (B) Reduction of CoV over the 6 training sessions. Asterisks indicate significant differences in variability between both groups at that specific training session (* ≤ .05, ** ≤ .001).

Fisher’s chi-square combination of rank-based partial *p*-values[51,54,55] across training sessions yielded a significant effect for group difference in performance variability (*p* = .01). Posthoc unadjusted and multiple testing adjusted comparisons revealed significant differences at TD1 (Brunner-Munzel *t*(34.20) = 2.16, *p* = .02, δ = .4 (medium)/ *p*FWE = .09) and TD4 (Brunner-Munzel *t*(39.94) = 3.45, *p* = .001, δ = .5 (large)/ *p*FWE = .03) displaying higher variation in the c-tDCS group performance than the s-tDCS group (Figure 4B). No significant group differences were found at TD2 (Brunner-Munzel *t*(34.76) = 0.95, *p* = .16/ *p*FWE = .40), TD3 (Brunner-Munzel *t*(41.99) = 1.66, *p* = .04/ *p*FWE = .17), TD5 (Brunner-Munzel *t*(41.69) = -0.85, *p* = .80/ *p*FWE = 0.81) and TD6 (Brunner-Munzel *t*(41.66) = 1.21, *p* = .12/ *p*FWE = 0.3)

### 3.3. Effect of concurrent tDCS on cognitive transfer

#### 3.3.1. Visual and Verbal Memory Test (VVM)

No effect of tDCS was found on delayed recall, Brunner-Munzel *t*(37.64) = -1.44, *p* = .16, δ = .25, or the rate of forgetting, Brunner-Munzel *t*(39.21) = 0.56, *p* = .58, δ = .1 (Supplementary materials 2.4.1, Supplementary Figure. S6).

#### 3.3.2. Trail making test (TMT)

ΔTMT. A noticeable improvement in ΔTMT was detected in the c-tDCS group compared to the s-tDCS group (Figure 5), Brunner-Munzel *t*(40.49) = 2.08, *p* = .04, δ = .34 (medium)

**Figure 5.**
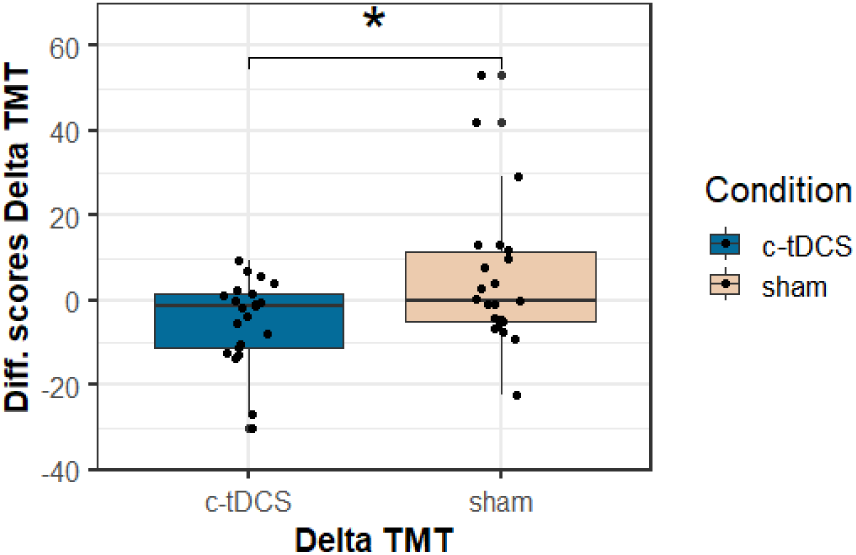
Delta TMT = TMT B – TMT A. Improvement in Delta TMT seen as pre-post difference scores calculated from the pre and post test scores expressed as seconds. Lower scores signify higher improvements. Asterisks indicate significant difference between groups.

TMT A. Both groups significantly improved in this subtest (Figure 6A) with no significant difference between either groups, Brunner-Munzel *t*(39.89) = -0.64, *p* = .52, δ = .12.

**Figure 6.**
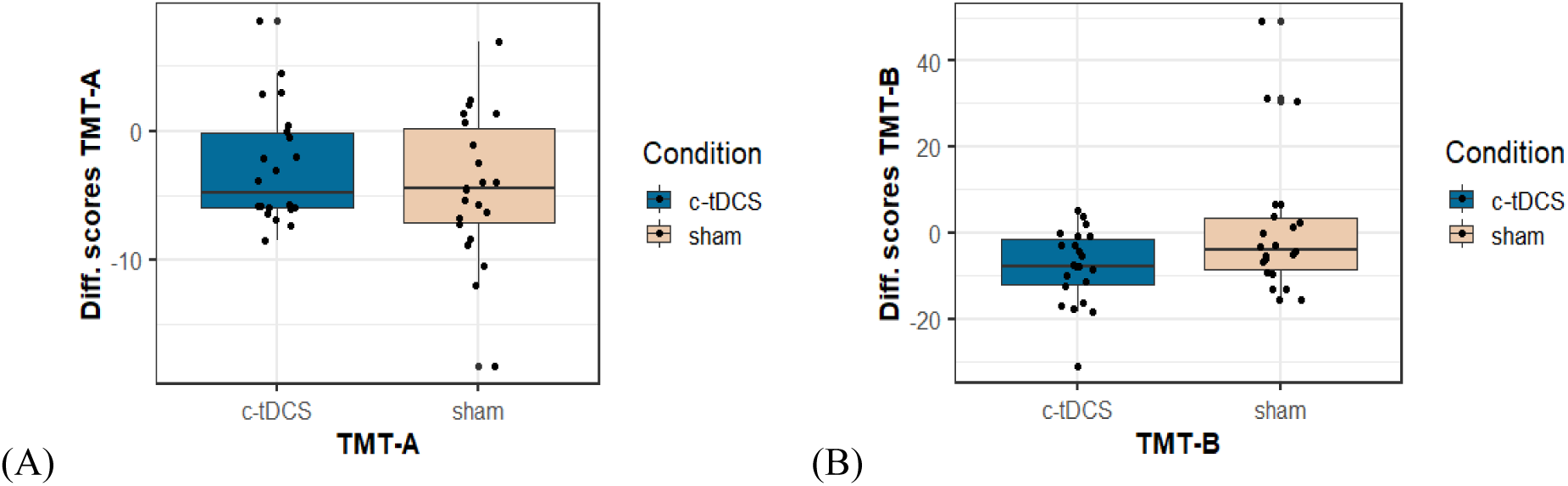
Trail-making test A & B: (A) Similar pre-post TMT-A difference scores for both groups; (B) Pre-post TMT-B difference scores for both groups displaying a tendency towards higher improvements in c-tDCS as compared to s-tDCS. Here lower scores signify good performance.

TMT B. In this subtest measuring cognitive flexibility, a trend towards higher improvements for the c-tDCS group compared to the s-tDCS group was observed (Figure 6B), Brunner-Munzel *t*(41.49) = 1.89, *p* = .07, δ = .31 (medium), implying faster completion times exhibited by the c-tDCS group compared to s-tDCS. Since the baseline performance in TMT-B was similar for both groups (Brunner-Munzel *t*(38.38) = -0.64, *p* = .53), justifications for such asymmetric performance improvement other than training under concurrent tDCS seem unlikely.

#### 3.3.3. D2-Test of Attention (d2)

Both groups improved in this test as observed in the concentration scores (Figure 7A), without significant difference between either groups(Figure 7 B), Brunner-Munzel *t*(40.67) = -0.99, *p* = .32, δ =

**Figure 7.**
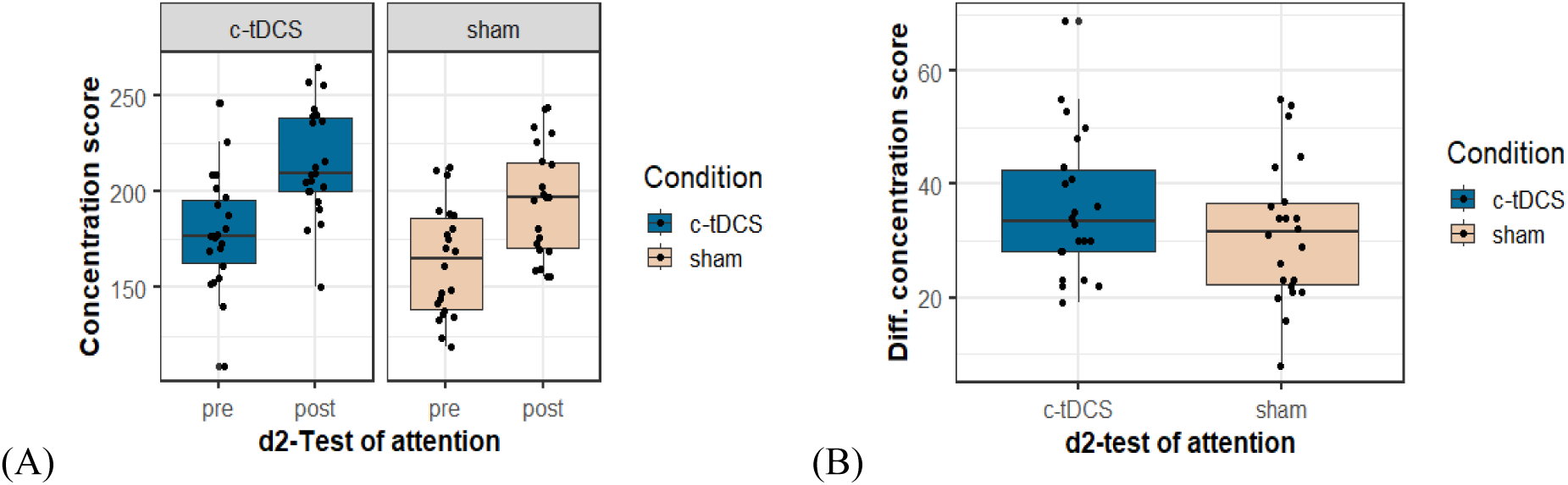
Concentration score was considered as a parameter of attention measured using d2-test of attention; (A) Pre- and post-test concentration scores for both groups; (B) Improvement in concentration scores seen as pre-post difference scores for both groups.

#### 3.3.4. Eriksen Flanker task

Accuracy interference. c-tDCS group showed comparatively lower improvements than the s-tDCS group in accuracy interference reduction after the intervention trending towards significance (Figure 8A), Brunner-Munzel *t*(41.83) = -1.93, *p* = .06, δ = .32 (medium)

**Figure 8.**
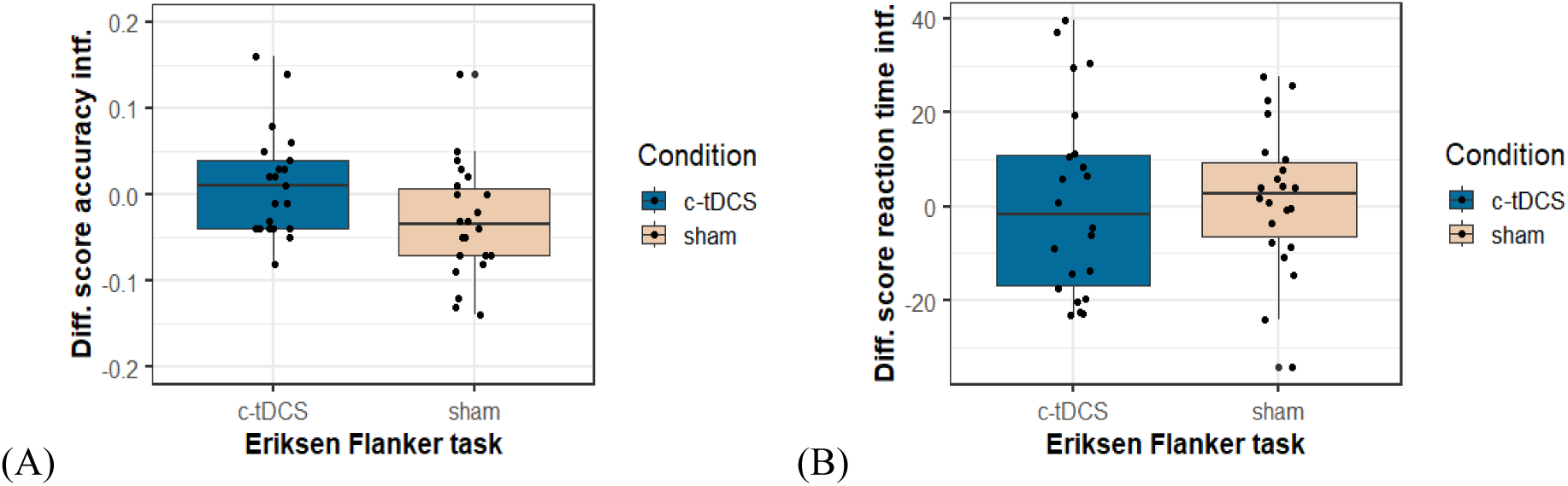
Accuracy interference and reaction time interference scores were considered as parameters of interest in the Eriksen flanker test: (A) Improvement in accuracy interference scores seen as pre-post difference scores calculated for both groups. For purposes of better visualization an outlier (-0.68) from the c-tDCS group was removed from the graph; (B) Improvement in reaction time interference scores seen as pre-post difference scores.

Reaction time interference. No difference between either groups was observed for this reaction time metric of the Eriksen flanker task, Brunner-Munzel *t*(35.40) = 0.31, *p* = .76 (Figure 8B).

### 3.4. Effect of intervention on motor transfer

Wii task. At the beginners level, both groups equally profited from the intervention, Brunner-Munzel *t*(40.34) = 0.15, *p* = .9, δ = .03 (Figure 9A). Whereas at the advanced level, although statistically not significant, c-tDCS group experienced higher improvements (median= 85 ± 76.6 points) than the s-tDCS group (median=61.7 ± 30.85 points), Brunner-Munzel *t*(28.75) = -0.79, *p* = .4, δ = .15 (Figure 9B).

**Figure 9.**
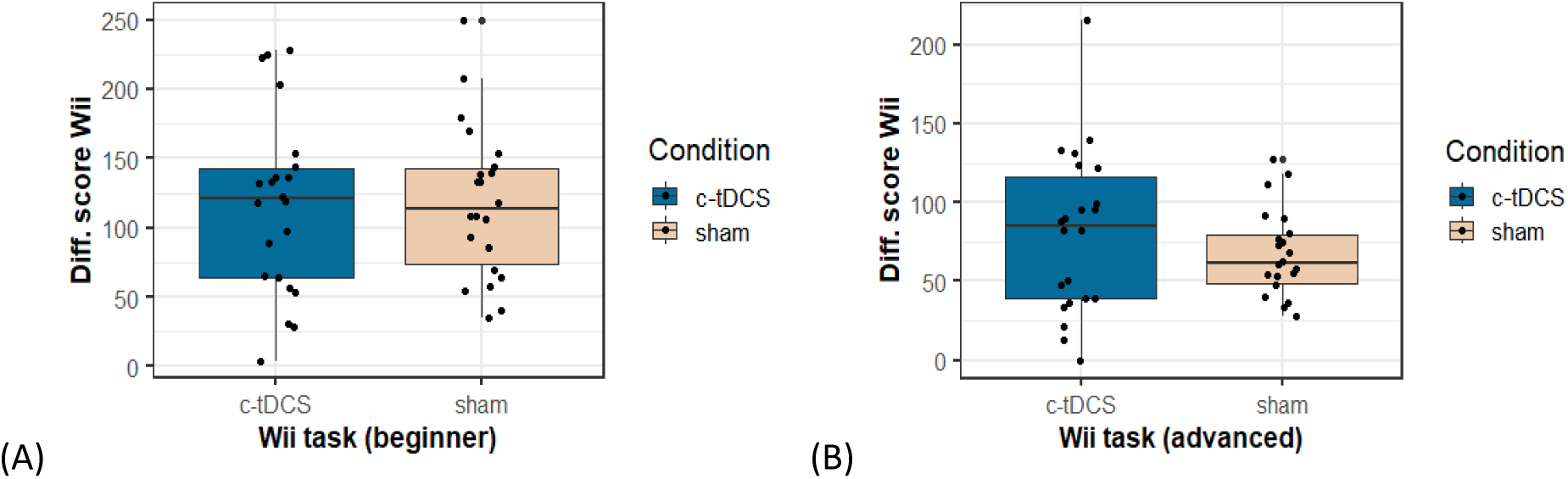
Effects of the interventions on performance of the football header task (Nintendo Wii). The Wii score is a cumulation of all the hits of the target objects and unsuccessfully dodged non-target objects; (A) Improvement in Wii scores at the beginner level; (B) Improvement in Wii scores at the advanced level are expressed as the difference in the pre to post test for both groups.

The difference scores revealed a clear distinction with-in the c-tDCS group where 13 participants exhibited larger improvements (≥80 points) in the Wii task (advanced level) compared to 9 participants with minimal gains (≤50 points). This distinction was absent in the s-tDCS group. A subsequent correlational analysis revealed a positive medium correlation between the difference scores of the Wii task (advanced level) and mean performance variability (CoV) over the entire training session on the stabilometer, which was only present in the c-tDCS group, *r* = 0.53, *p* = .01, not observed in the s-tDCS group (*r* = -0.20, *p* = .39). This suggests that participants from the c-tDCS group exhibiting larger variability during stabilometer practice also displayed higher gains in the Wii task (Figure 10).

**Figure 10.**
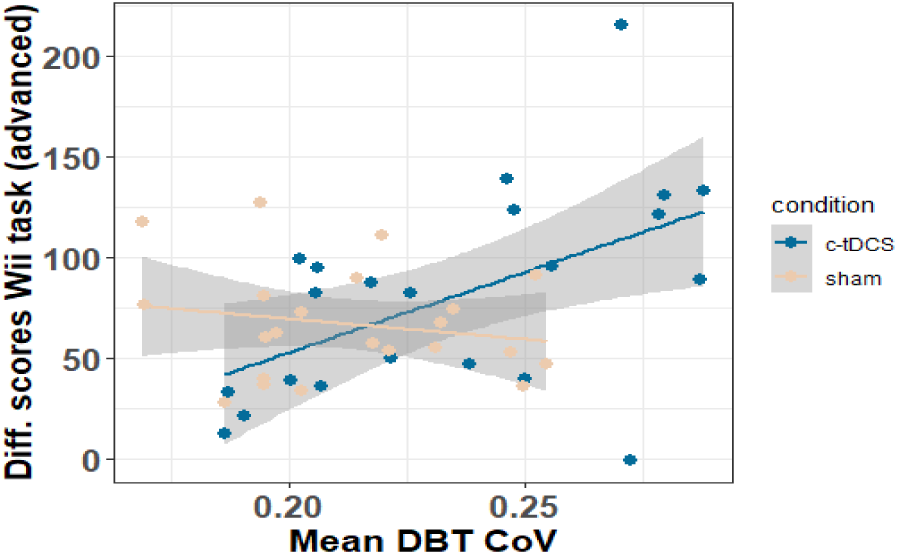
Correlation between performance variability (CoV) on the stabilometer and Wii change scores (advanced level; pre to post intervention) for the c-tDCS and s-tDCS groups.

## 4. Discussion

This randomised, double-blinded, sham-controlled tDCS study highlights the importance of frontal networks in learning a complex dynamic balance task. Our results demonstrate that the influence of c-tDCS over these networks during a long-term motor learning process caused higher performance variability compared to the s-tDCS stimulation group. This increase in behavioural variance indicates that the stimulation causally affected (pre-)frontal brain networks [27,28]. Moreover, DBT training with concurrent c-tDCS not only resulted in a ‘near’ transfer effect on postural control, but also in ‘far’ transfer on cognitive flexibility known to rely on the prefrontal networks persisting 24 hours after the end of training.

### 4.1. PFC involvement in balance learning

In this study, tDCS applied during DBT practice was aimed at influencing network nodes implicated in long-term DBT learning. Hence, shifting the focus onto the specific task-relevant activation of networks, down-weighing the low anatomical precision of tDCS[28,59]. These network nodes were selected based on previous findings showing macro- and microstructural properties of PFC-SMA regions predict future DBT learning[10,60] also changing in response to DBT practice[11,61,62]. Although these studies provided evidence of a brain-behaviour relationship between PFC-SMA networks and balance learning, demonstrated using approaches like statistical mediation analyses [63], the neuroimaging findings remain correlative. However, [35] showed a single session of online c-tDCS over the right PFC-SMA region during training has an acute effect on subsequent DBT performance. Here, we extend these previous findings by causally showing PFC-SMA network involvement in long-term balance learning, manifesting itself through increased performance variability [28].

The true direction of the effect of tDCS on performance may be masked/varied across and with-in participants due to dissimilar amplification in neuronal noise, in such cases, the sheer increase in variance (beyond measurement noise) after tDCS may be considered evidence for a cause–effect relationship[28]. Such behavioural consequences of tDCS may arise due to individual differences in the recruitment of brain networks during task performance leading to differences in excitability modulation [20,28,64]. Along with reported within-session, non-linear effects of c-tDCS[65], dissimilarities in tDCS induced modulation of cortical excitability may not necessarily translate into behavioural deviations as drastic as performance inhibition. Lack of DBT performance deterioration can therefore be associated with tDCS being a weak direct current and its behavioural effects meagre; making it possible for networks to capably compensate for weak disturbances during online stimulation by adapting to the electric field over time[28]. The results of this study demonstrate improved DBT performance for both groups over the 3-week training duration; indicating similar task proficiency at the end of practice. Hence, tDCS may have affected the process of learning a complex task rather than altogether changing the learning trajectory.

The prefrontal networks involved in the strategy building aspect of motor learning were the prime target of c-tDCS in our study [24,59]. Consequently, participants were not instructed on the most optimal task execution strategy (contrary to a ‘classical’ motor skill learning/training), instead, encouraged to learn the task by discovering their own strategies via trial and error[38]. Previous studies investigating the mechanisms involved in adopting specific courses of action during learning have associated the anterior PFC in exploration of new possibilities. Here, future outcomes are said to be predicted by tracking alternative options and exploratory switching between courses of actions through extrapolation of short-term trends [7,9]. Hence, task complexity and uncertainty of outcomes may dictate the extent of PFC involvement, where selection of appropriate strategies and guiding cognitive resources to implement these strategies is done by integrating and comparing various sequential outcomes [6,9]. Owing to the task complexity and the available solution space, the DBT fulfils criteria’s particularly conducive for cognitive processes involved in reinforcement learning, in particular, exploration of solutions achieved through various coordinative whole-body movements. Therefore, we speculate that PFC-dependent networks responsible for exploration of new performance strategies (in the context of learning) were modulated by c-tDCS. This modulation was behaviourally expressed as increased performance variability.

### 4.2. PFC and balance training-induced transfer

It is suggested that extending learning gains to other untrained tasks is possible only if a shared commonality exists between these tasks, viz., abilities required in executing both tasks, neural processing mechanisms and brain regions [16,17,66]. These transfer effects are also theorised to be tied to early phases of structural plasticity within overlapping networks[19]. The ‘neural overlap hypothesis’ has been supported by evidence from concurrent tDCS during cognitive training resulting in microstructural brain alterations alongside near-transfer behavioural effects [34,67]. Since the motor learning paradigm used in this study is capable of inducing structural grey and white matter changes in PFC and SMA regions [11,62,68,69], we further hypothesized it to potentially lead to cognitive transfer effects. Consistent with this hypothesis, we found higher improvement in executive functioning performance (i.e., ΔTMT and TMT-B)[70] as a result of DBT training with concurrent rPFC c-tDCS compared to s-tDCS. Both, aerobic exercise on its own[71] and a-tDCS over left DLPFC during coordinative exercise [44] have shown a tendency towards TMT performance improvements. Similarly, cognitive training combined with tDCS at an intensity of 1.0-mA augmented both decision-making performance and cognitive transfer[72].

Despite a global network involvement in TMT execution[73], our regions of interest were restricted to the overlapping PFC-SMA networks involved in DBT learning. We hypothesize the combination of DBT training-induced plasticity, discovery-learning based motor training and tDCS to encourage a rapid network reorganisation and compensation [74–76]. This functional compensation probably constituted conditioning new or otherwise inactive networks within the overlapping brain regions leading to an advantageous effect of intervention, absent in the s-tDCS group [77,78]. Benefiting from richly connected brain networks supporting a multitude of cognitive functions required in TMT-B execution may have improved the potential for transfer via compensatory mechanisms in the overlapping networks [73,79–81]. A combination of brain imaging and stimulation techniques is required to prove the specific functional and structural correlates of PFC involvement in learning and associated transfer.

Contrary to executive functioning, we did not find significant differences between either groups on memory and attention abilities, although positive effects of physical exercise (e.g., coordinative and aerobic exercise) on visuospatial attention, working memory [82], associative memory, spatial cognition [14,15] and visuospatial memory [83] have been observed in previous studies. Note, however, that our results indicate marginally better performance in the attention task (d2-R) exhibited by the c-tDCS group compared to the s-tDCS group. Although this difference did not reach statistical significance. On the other hand, the s-tDCS group showed a tendency towards higher improvements in an SMA-dependent selective interference resolution task (Eriksen flanker task-accuracy interference) as compared to the c-tDCS group, this trend was accompanied by a medium sized effect (Results 3.3.4).

Finally, the observed transfer effects on PFC-SMA-dependent cognitive tasks can be assumed to be due to a shared commonality with the trained task (neural overlap hypothesis)[19,66], which changed as a function of the intervention, demonstrating a potential common neural substrate underlying the trained balance task and the transfer task[84]. This complex motor training engaging higher-order processes may have enabled cognitive improvements by transferring learning gains to untrained tasks. In turn benefiting abilities like information processing, goal-dependent inhibition/ maintenance of responses, formulating strategies based on feedback, distributing attention over multiple strategies, switching between strategies(cognitive flexibility), etc [16,17,66]. Findings from [14] demonstrate balance training-induced improvement in memory and spatial cognition attributed to a training that encompassed proprioceptive, visual and motor-based learning. Likewise, a month of slackline training improved vestibular-dependent spatial orientation performance [13] suggesting a positive effect on vestibulo-hippocampal spatial orientation.

Lastly, we also observed a statistical tendency towards larger near-transfer effects to an untrained balance task (Nintendo Wii header game-advanced level) in the c-tDCS group compared to the s-tDCS group. Interestingly, consistent with the ‘neural overlap hypothesis’, in the c-tDCS but not in the s-tDCS group we observed a medium-sized positive correlation between DBT performance variability and Wii scores. Such near motor transfer effects have recently been observed by [85], manifested as improved cross-limb transfer from the trained to the untrained hand after anodal tDCS over rM1 in older adults. Similarly, we hypothesize that participants in our study were able to successfully use the movement solutions learned during DBT training onto an untrained balance task which also requires a comparable movement pattern in terms of body’s centre of mass (COM) control and displacement. [86–88] emphasize introduction of variation during practice as a key aspect in eliciting new movement solutions enabling a degree of transfer beyond the practiced solutions. However, further studies are required to support the role of movement variability to improve transfer during stabilometer learning.

### 4.3. Limitations

Although the results of this study highlight the importance of the frontal networks in learning a complex task, we are unable to disentangle the contributions of PFC from those of SMA as both these regions have been implicated with undergoing learning-induced structural changes. Our cognitive transfer results do point towards higher PFC involvement but we were not able to definitively outline the specific contributions of these regions. The utilization of a combination of tDCS and neuroimaging may aid in explicitly mapping stimulation-induced changes at the neuronal and network levels. Linking these brain changes to the behavioural effects would be the natural subsequent step in order to unravel the complexity of the underlying brain-behaviour relationship. Stimulating an alternative brain region is advised in order to ascertain that the observed effects emanate solely as a result of interference within the regions of interest [28,29]. However, this control condition was not included since we intended on influencing the networks previously implicated in learning the complex DBT. In light of the recently revealed predispositions to improved learning abilities [10,60], heterogeneity of participants in the form of genetic makeup, brain structure and environmental diversity requires consideration [89]. The solitary effect of tDCS on cognitive abilities without the influence of training is an aspect that could help differentiate between the cumulative effect of tDCS and training observed in this study.

## 5. Conclusion

Our results provide new evidence for PFC-SMA involvement during long-term DBT practice. Specifically, we show that interfering with these networks using c-tDCS leads to increased performance variability, potentially indicating a causal involvement of PFC-SMA networks in DBT learning[28]. Against the background of ‘neural overlap hypothesis’, we interpret the observed tDCS-effects on motor and cognitive performance as tDCS effects pertaining not only to the trained tasks, but also to the untrained tasks which rely on overlapping brain networks. The conclusions drawn through this study reinforce the positive impact of physical activity on cognition through the synergistic neural networks sub-serving both motor processing and cognitive functioning. An understanding of this brain-behaviour relationship may prove valuable not only in promoting overall health through exercise but also support healthy aging by means of mobilizing neural resources to remedy dysfunction.

## CRediT author contributions

NP: Conceptualization, methodology, data acquisition, formal analyses, data curation and writing-original draft, review & editing;

NL: Conceptualization, methodology, supervision, formal analyses and writing-review & editing; EK: Methodology, formal analyses, writing-review & editing;

NM: Methodology, resources, writing-review & editing;

MT: Conceptualization, methodology, resources, funding acquisition, supervision, formal analyses and writing-original draft, review & editing

## Acknowledgements

The authors would like to thank Fredericke Schökel for her help with German translations of study documents and assisting in data acquisition. We would also like to thank Lukas Hennecke for assisting in data acquisition and Marlene Schmicker & Inga Menze for their expert opinion and guidance related to tDCS.

