## Supplementary material for "Online stimulation of prefrontal cortex during practice increases motor variability and modulates later cognitive transfer: a randomized, double-blinded & sham-controlled tDCS study"

#### 1. Methods

1.1. Sample size estimation: Sample size estimation for this study was based on the expected behavioural effect of tDCS on online DBT performance. In this respect, [1] revealed the sham group to significantly outperform the interfering stimulation group in terms of online learning performance with a large underlying effect size ($d \approx 0.9$, as estimated from the reported F-statistic). Assuming a slightly lower effect size of $d = 0.8$, probability of type I error $\alpha = 0.05$, and power = 0.8 yielded a sample size of n = 21 per group (two-sample t-test, one-tailed; [2]).

1.2. Transfer tasks:

1.2.1. Visual and Verbal Memory Test (VVM; Schächtele & Schellig, 2009): evaluates short-term maintenance of memory consisting of two subtests assessing visuospatial memory and verbal memory separately. Only the visuospatial subtest represented by a street map was used in this study as numerous studies have shown coordinative training [4,5], aerobic training [6] as well as tDCS-based [7] interventions to influence working memory, in particular visuospatial memory functions. This subtest requires the participants to memorise a given path on a street map and recall it (immediate and delayed) on an identical but empty street map. Parallel forms of the subtest (theatre & museum) were employed for pre- and post-testing. Encoding time was fixed at 2 minutes shortly followed by time for retrieval (free immediate recall-VVM1) timed at 2 minutes to completion while delayed recall (VVM2) of the material was performed 30 minutes later. The performance measures of interest were the number of correctly recalled intersections in the street map, with a maximum of 31 correct intersections, and rate of forgetting calculated as a percentage of the immediate and delayed recall using:

$$\text{Rate of forgetting} = (\text{VVM2} - \text{VVM1}) / \text{VVM1} * 100$$

1.2.2. D2- Test of Attention: assesses sustained attention and visual scanning speed and accuracy [8]. Findings from [9] demonstrated cardiovascular exercise-induced frontotemporal plasticity to mediate improved attention measured using this test, also assumed to be an ability vital in learning the DBT. We used the German paper and pencil version of the revised “d2 Test of Attention” (d2-R; [10]) which consists of 14 test lines with 47 symbols per line. The symbols can be either of the lowercase letters “d” or “p” marked with 1, 2, 3, or 4 small dashes above and/or below the letter. The participant is required to strike through occurrences of the letter “d” bearing 2 dashes only as quickly and accurately as possible. All other symbols act as distractors to be ignored. The participants were requested to proceed from left to right with 20 seconds dedicated to each line; after which they need to proceed to the next line. Concentration performance was the variable of interest, defined as the number of marked distractors (sum of errors of commission and errors of overlooking) deducted from the total number of processed targets.

1.2.3. Eriksen Flanker task [11]: addresses interference resolution ability, a more selective inhibition process, where task-relevant responses are maintained while task-irrelevant stimuli and goal-irrelevant responses are inhibited [12,13]. Exercise interventions have not only shown to positively impact this interference resolution ability [14,15], but the neural correlates of this task also correspond with areas previously shown to undergo structural changes after DBT learning [12,16,17]. In this task, the participants were presented with a fixation arrow at the centre of the screen which was then replaced by a stimulus cue (> or <). Here they were asked to respond only to the stimulus cue not the flanking array of arrows on either side. The trials consisted of either flanking arrows pointing in the same direction as the central stimulus cue (congruent trial: < < < < <), or flanking arrows pointing in the opposite direction (incongruent trials: > > < > >). Each participant underwent a familiarisation block of 10 trials followed by two successive blocks with 50 trials each, first-order counterbalanced such that congruent and incongruent trials followed each other equally as often. Keeping the target stimulus as simple as possible with ‘<’ representing left button press and ‘>’ representing right button press avoided incorrect responses due to failure of learning the correct responses; hence focusing on target identification, information processing followed by the effect of response selection conflict [11].

1.2.4. Trail making test (TMT; [18,19]): is a measure of divided attention and scanning abilities, with particular focus on cognitive flexibility involving switching between sets of letters and alphabets. The neural correlates of this task also largely overlap with the areas of interest in this study [20–22]. This test was conducted in 2-parts: (i) part A requires the participant to connect randomly scattered numbers from 1 to 25 in an ascending order; (ii) part B requires the participant to alternate between letters (A-L) and numbers (1-13), i.e., connecting a number to an alphabet moving in an ascending order (for numbers) and alphabetical order, constantly switching between these sets (1-A-2-B-3-C). The participant is required to familiarise themselves with shortened versions of each part before they begin. They are instructed to perform the task as fast as possible with minimal errors without lifting the pen off the sheet. Time to completion and difference between the time required to complete parts A & B (delta/Δ TMT) are used as variables of interest.

1.2.5. Wii task (Motor transfer test): A football header task available on the WiiFit console was used to assess the goal-directed control of COM movement/sway of the participants (accuracy as well as reaction time), similar to movement control required during execution of the DBT. A Wii Nintendo connected to a large TV screen was used for this task. The participants were asked to stand with both feet on the Wii-board where their centre of mass was tracked and represented as an avatar on the screen with the goal of heading balls kicked in their direction in order to gain points. Simultaneously, other objects (panda masks and shoes) were also added to the mix with instructions to dodge these objects or risk losing a point. These objects were tossed either laterally (left or right) or in the centre, prompting the participant to accurately realign the avatar in the required direction by shifting their COM/ body weight on the Wii board. Participants performed 10 trials with 30sec breaks in between. The total score, a measure of both accuracy and reaction time, is derived through the successful hits and misses of objects during each trial.

1.3. Control tasks:

1.3.1. Balance Error Scoring System (BESS) [23]: This test includes maintaining various stances/positions executed bare feet, hands on the hips, eyes closed, initially on the floor followed by balancing on a medium density foam cushion. The different test positions comprised: double leg stance, single leg stance and a tandem stance. During testing, the participants were asked to maintain the test position for a duration of 20secs each with as few deviations as possible. Every deviation from the original position was marked as an error, with a maximum score of 60. As prescribed in the BESS protocol, errors included opening of eyes, stepping, moving out of position, hands leaving hips, etc. The summed-up error scores from all the test positions in the pre and post-test at all three training sessions were used for analyses.

1.3.2. Stroop test [24,25]: To control for the effect of tDCS on general executive functions, a computerised version of Stroop test was administered using the Presentation® software by Neurobehavioral Systems, Inc., Berkeley, CA, USA. Coloured words reading red, blue, green or yellow were presented on a screen with a white background either in a congruent or incongruent font colour. Buttons on the keyboard pertaining to the respective colours were pre-assigned. The task included two conditions, viz., participants asked to respond by identifying either the colour of the word or the word itself by pressing the pre-assigned button as soon as possible. Prior to testing, all participants went through familiarization trials. Accuracy as well as quick response were emphasized as good performance. The participants received immediate feedback after every trial as ‘correct’ or ‘incorrect’ response. A total of forty-eight trials per condition were presented in a randomized order (congruent/incongruent). For the purpose of analyses, an interference score was calculated (outcome measure: mean reaction time or accuracy for the incongruent condition subtracted from the congruent condition)

### 2. Further analyses and results:

2.1. Stimulation questionnaire: Mean intensity of the sensations reported by subjects after tDCS (3.1.1). Intensities for tingling, burning, headache, nausea and ill feeling were evaluated on a 5-point scale with 1: none at all, 2: mild, 3: moderate, 4: somewhat strong/ considerable, 5: very strong. The effect of tDCS on concentration, attention, alertness, mood change was rated on an inverted scale with 1: strong disturbance, 2: mild disturbance, 3: no effect, 4: mildly improved, 5: strongly improved (Supplementary Table S1).

*Supplementary Table S1. Stimulation questionnaire responses for subjective perception of tDCS immediate effects. Values denote mean rating for each sensory perception separated for each group.*

| **Training session** | **Condition** | **Tingling** | **Burning** | **Headache** | **Nausea** | **Feel ill** | **Concentration** | **Attention** | **Alertness** | **Mood change** |
|---|---|---|---|---|---|---|---|---|---|---|
| **1** | s-tDCS | 2.22 | 1.61 | 1.52 | 1.05 | 1.24 | 1.52 | 3.01 | 3.08 | 3.05 |
| **1** | c-tDCS | 2.24 | 1.58 | 1.46 | 1.04 | 1.24 | 1.50 | 3.01 | 3.08 | 3.06 |
| **3** | s-tDCS | 2.25 | 1.62 | 1.52 | 1.05 | 1.25 | 1.54 | 3.00 | 3.07 | 3.05 |
| **3** | c-tDCS | 2.26 | 1.59 | 1.46 | 1.03 | 1.23 | 1.50 | 3.02 | 3.09 | 3.06 |
| **5** | s-tDCS | 2.27 | 1.62 | 1.52 | 1.05 | 1.25 | 1.52 | 3.01 | 3.06 | 3.04 |
| **5** | c-tDCS | 2.24 | 1.59 | 1.46 | 1.03 | 1.23 | 1.48 | 3.04 | 3.11 | 3.06 |

2.2. Baseline performance: Baseline performance recorded as the first 2 trials on TD1 (before tDCS stimulation commenced) was found to be similar between both groups (mean TIB c-tDCS: 3.05 ± 1.7 secs vs s-tDCS: 2.99 ± 1.49 secs), Brunner-Munzel $t(41.97) = 0.27$, $p = .80$, Cliff δ = 0.04 (Supplementary Figure S1).

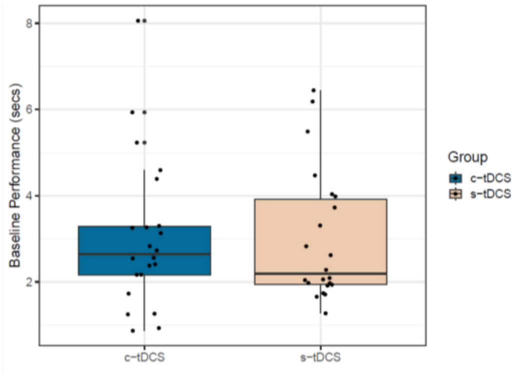


*Supplementary Figure S1. Baseline performance on the DBT recorded at the first 2 trials (without tDCS stimulation), was found to be similar between both groups. Y-axis represents the TIB in seconds.*

2.3. Minimum values. A comparison of the minimum (lowest TIB) and maximum (highest TIB) values attained by every participant during the entire training duration revealed a progressive improvement in the minimum values over the training period $F(5, 19.13) = 59.35$, $p = .00$, $d>1$ , along with an effect of interaction $F(5,19.13) = 3.45$, $p = .02$, $d = 0.57$ without a group effect $F(1, 25.49) = 0.42$, $p = .5$ (Supplementary Figure S2). The maximum values did not exhibit any effects of interaction or group, instead only an effect of time was detected $F(1, 19.11) = 18.86$, $p = .00$, $d>1$.

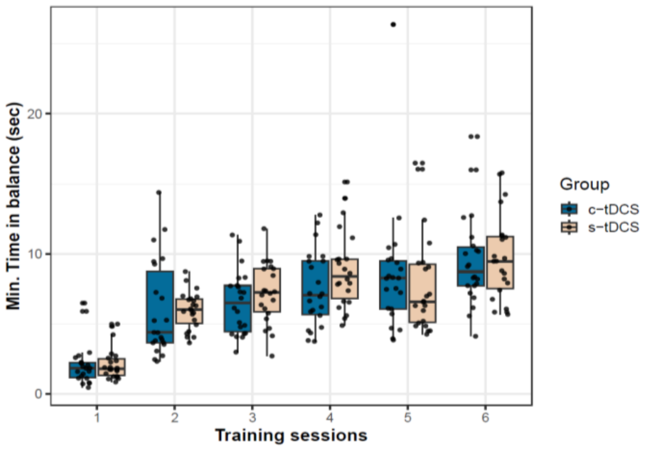


*Supplementary Figure S2. Minimum TIB values exhibited by every participant in both groups over 6-training sessions.*

2.4. Cognitive and control tests:

2.4.1. VVM: No differences between groups were found in this visuospatial memory test (Supplementary figure S3 and 3.3.1 in manuscript)

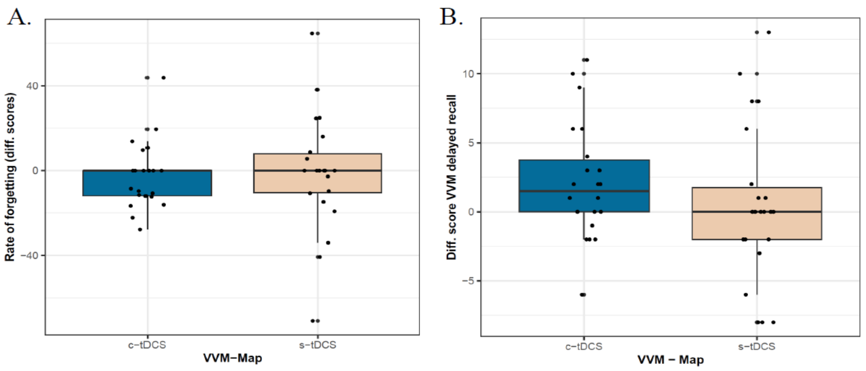


*Supplementary Figure S3. Improvement in visuospatial memory task VVM seen as pre-post difference scores calculated from the pre and post test scores for (A) rate of forgetting and (B) delayed recall.*

2.4.2. TMT-B: A negative correlation only in the s-tDCS group was found between the difference scores of the TMT-B subtest and the mean CoV (Supplementary Figure S4) with a trend towards significance ($p$= .06, $r$ = - 0.41) suggesting that participants with lower CoV did not improve in this task as much as the participants exhibiting higher variability.

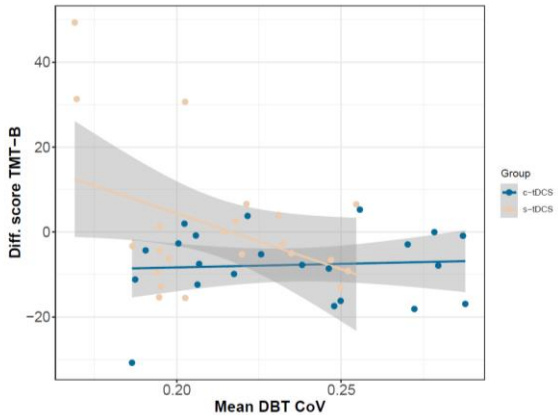


*Supplementary Figure S4. Correlation between motor variability (CoV) on the stabilometer and the difference score in TMT-B subtest in the c-tDCS and s-tDCS group.*

2.4.3. Eriksen flanker task: A significant positive correlation as seen in Supplementary figure S5 was noticed between the accuracy interference score and the mean CoV only in the s-tDCS group ($p$

= .05, *r* = 0.43) implying participants with a lower variability experienced higher gains in accuracy by successfully reducing their accuracy interference.

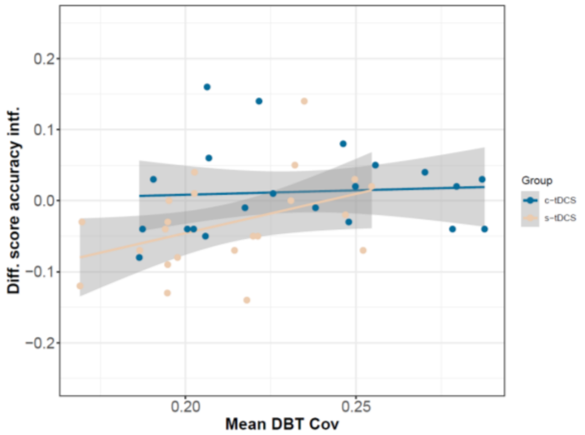


*Supplementary Figure S5. Correlation between motor variability (CoV) on the stabilometer and the difference score in accuracy interference values in the c-tDCS and s-tDCS group. For purposes of better visualization an outlier with a score of (-0.68) from the c-tDCS group was removed from the graph.*

2.4.4. BESS score: No differences between groups were observed for this control measure of general balance ability (Supplementary figure S6 and 3.1.4 in manuscript).

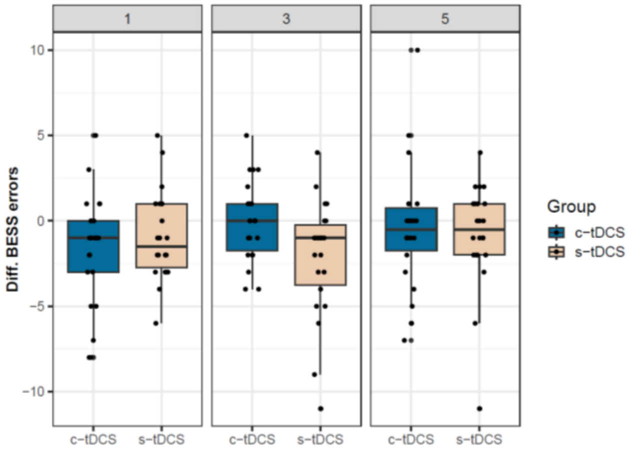


*Supplementary Figure S6. Improvement in general balance ability measured using BESS. Y-axis shows pre-post difference scores calculated immediately before and after intervention on TD1, TD3, TD5.*

2.4.5. Stroop task: No difference was observed between groups in the accuracy interference part of the Stroop test, however in the reaction time part of this test, an effect of group without an interaction effect or time was observed (Supplementary figure S7 and 3.1.3 in manuscript).

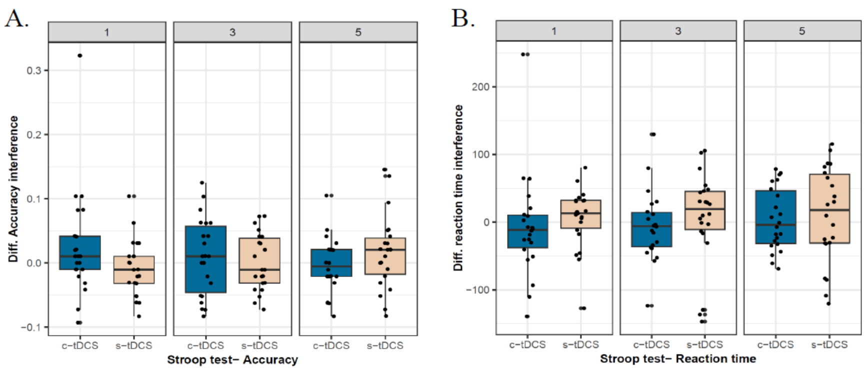

*Supplementary Figure S7. Changes in general cognitive ability measured using the Stroop task. Y-axis shows pre-post difference scores calculated immediately before and after intervention on TD1, TD3, TD5 (A) accuracy interference= difference between incongruent and congruent responses; (B) reaction time interference= difference between reaction time of incongruent and congruent responses.*
